# Recombination and purifying selection preserves covariant movements of mosaic SARS-CoV-2 protein S

**DOI:** 10.1101/2020.03.30.015685

**Authors:** Massimiliano S. Tagliamonte, Nabil Abid, David A. Ostrov, Giovanni Chillemi, Sergei L. Kosakovsky Pond, Marco Salemi, Carla Mavian

## Abstract

In depth evolutionary and structural analyses of severe acute respiratory syndrome coronavirus 2 (SARS-CoV-2) isolated from bats, pangolins, and humans are necessary to assess the role of natural selection and recombination in the emergence of the current pandemic strain. The SARS-CoV-2 S glycoprotein unique features have been associated with efficient viral spread in the human population. Phylogeny-based and genetic algorithm methods clearly show that recombination events between viral progenitors infecting animal hosts led to a mosaic structure in the S gene. We identified recombination coldspots in the S glycoprotein and strong purifying selection. Moreover, although there is little evidence of diversifying positive selection during host-switching, structural analysis suggests that some of the residues emerged along the ancestral lineage of current pandemic strains may contribute to enhanced ability to infect human cells. Interestingly, recombination did not affect the long-range covariant movements of SARS-CoV-2 S glycoprotein monomer in pre-fusion conformation but, on the contrary, could contribute to the observed overall viral efficiency. Our dynamic simulations revealed that the movements between the host cell receptor binding domain (RBD) and the novel furin-like cleavage site are correlated. We identified threonine 333 (under purifying selection), at the beginning of the RBD, as the hinge of the opening/closing mechanism of the SARS-CoV-2 S glycoprotein monomer functional to hACE2 binding. Our findings support a scenario where ancestral recombination and fixation of amino acid residues in the RBD of the S glycoprotein generated a virus with unique features, capable of extremely efficient infection of the human host.

## Introduction

Coronaviruses (CoVs) are single strand, positive sense single-stranded RNA (+ssRNA) viruses, with diverse tropism, able to infect respiratory, enteric, and hepatic tissues of several species (Fehr and Perlman 2015). Within the CoVs family, Beta CoVs have repeatedly proven the ability to shift from their natural reservoir and adapt to human hosts (Su, et al. 2016). Several CoVs, such as HCoV-229E, HCoV-OC43, HCoV-HKU1, and HcoV-NL63, are mostly associated with mild symptoms (Bucknall, et al. 1972; Woo, et al. 2005). Few CoVs caused limited outbreaks of more severe illness, such as the severe acute respiratory syndrome (SARS) and Middle East respiratory syndrome (MERS) (Leung, et al. 2004; Zaki, et al. 2012), and the recently emerged SARS-CoV-2 (Wu, et al. 2020)). First reported in China in December 2019, CoV disease 2019 (COVID-19) quickly turned into a pandemic, with nearly 7,000,000 cases and hundreds of thousands of deaths reported worldwide as of June 2020 (Kamel Boulos and Geraghty 2020). An early study hypothesized SARS-CoV-2 emergence from snakes (Ji, et al. 2020), but further phylogenetic analyses traced its origins to CoVs circulating in bat (Zhou, Chen, et al. 2020; Zhou, Yang, et al. 2020) and pangolin (Xiao, et al. 2020). Contacts between different animal species prior to human exposure might facilitate host shift, possibly through adaptation to intermediate hosts, such as camels in the case of MERS-CoV (Dudas and Rambaut 2016; Su, et al. 2016) or civets or pigs in the case of SARS-CoV-1 (Wang and Eaton 2007).

While most RNA viruses use low-fidelity RNA polymerases, CoV genomes encode a proofreading and mismatch correction pathway, resulting in less error prone replication and translating into lower mutation rates (Ferron, et al. 2018; Li, Wang, et al. 2020). This feature might ultimately limit the mutagenic variability of the virus, and recombination might prove a prominent driving factor in the evolution and expansion of species range and cellular tropism. Recombination can take place whenever a single host is infected with multiple, genetically different, strains, and is an important factor driving viral diversity (Simon-Loriere and Holmes 2011; Mavian, et al. 2017). Recombination has been reported as a possible mechanism favoring cross-species transmission of CoVs and increasing their adaptability to new host (Graham and Baric 2010; Su, et al. 2016). There is copious evidence of such events having occurred in the ancestral history of both MERS-CoV and SARS-CoV-1 (Lau, et al. 2015; Dudas and Rambaut 2016; Anthony, et al. 2017) and CoVs in general (Lai, et al. 1985; Decaro, et al. 2009; Terada, et al. 2014; Tian, et al. 2014), but evidence regarding the recombinant origin of SARS-CoV-2 is not conclusive (Ji, et al. 2020; Liu, et al. 2020; Paraskevis, et al. 2020; Zhou, Chen, et al. 2020). Paraskevis *et al*., who made the case for a non-recombinant origin of the virus, used a limited dataset, which did not include pangolin CoVs (Paraskevis, et al. 2020). Based on genetic similarity plots, Ji *et al*. and Liu *et al*. hypothesized a possible recombination event with an undetermined viral isolate in the ancestry of SARS-CoV-2 (Ji, et al. 2020; Liu, et al. 2020). A very recent publication by Li *et al*. (Li, Giorgi, Marichannegowda, et al. 2020) also hypothesized a recombination event between bat and pangolin viruses as progenitor of the human CoV-2. While recent studies have used simple recombination detection algorithms, leading to various results that may depend on the assumptions of the specific algorithm and the data set of choice (Boni, et al. 2020; Li, Giorgi, Marichann, et al. 2020; Wong, et al. 2020), we performed an in-depth analysis of SARS-CoV-2 and closely related SARS-CoV-1 and SARS-like CoVs lineages using seven different methods, including phylogeny-based and genetic algorithms (see Methods section) (Kosakovsky Pond, et al. 2006; Martin, et al. 2015), with the purpose of achieving higher sensitivity (Jia, et al. 2018). Our analysis in fact revealed that the recombination pattern of the S glycoprotein is far more complex than previously thought. Because of the lack of sensitive recombination methods (Li, Giorgi, Marichannegowda, et al. 2020) and the comparison with only close relative of SARS-CoV-2, i.e. bat-SARS-CoV-2 RaTG13 and RmYN02 (Zhou, Chen, et al. 2020), it is possible that the likely mosaic nature of the SARS-CoV-2 genome was masked.

The SARS-CoV-2 S glycoprotein trimer is responsible for the crucial step of fusion of the viral membrane with the host cell membrane (Gallagher and Buchmeier 2001; Simmons, et al. 2013). Complex structural rearrangements follow the interaction of RBD (that must be in an exposed state referred as “up” conformation) with the peptidase domain (PD) of hACE2 (Wrapp, et al. 2020; Yan, et al. 2020). Conversely, the “down” conformation corresponds to the receptor-inaccessible state. The RBD up/down rotation is made possible by a hinge region at the N-term of RBD (Wrapp, Wang et al. 2020) and our Molecular Dynamics (MD) simulations identified threonine 333 as the hinge residue. A two-step sequential protease cleavage model has been proposed for activation of CoV S proteins, as shown for SARS-CoV and MERS-CoV (Belouzard, et al. 2009; Millet and Whittaker 2014), priming cleavage at the S_1_/S_2_ domain junction (Li 2016; Gui, et al. 2017), allowing the projection of the fusion peptide toward the target cell membrane (Bosch, et al. 2008) and activating cleavage on S2’ site (Ou, et al. 2020).

An important peculiarity of SARS-CoV-2 as compared to other CoVs is the presence of a furin-like cleavage sequence (Madu, et al. 2009; Lai, et al. 2017). Exposure of this cleavage site to the soluble furin protease is an important step in the viral infection (Wrapp, et al. 2020), and explain, at least in part, the enhanced infectivity and pathogenicity of the virus (Bosch, et al. 2003).

## Results

### On the origin of SARS-CoV-2 lineage: recombination in bat and pangolin

Since the human CoV isolates are ~96% identical to bat-SARS-CoV-2 (Andersen, et al. 2020), but ACE2 binding residues F486, Q493, S494 and N501 (Figure 1c) are shared with pangolin SARS-CoV-2 lineage b isolates, we investigated the evolutionary relationships among available full genome sequences (bat and pangolin-related SARS-CoV-2 isolates, bat-SL-CoVs, SARS-CoV-1, and SARS-like isolates) using a neighbor network (NNet) based algorithm (Bryant and Moulton 2004). Unlike standard phylogenetic inference methods, network approaches can naturally represent the presence of conflicting phylogenetic signal in the data (which is a hallmark of recombination), as we found in the analysis of the coronavirus genomes (Figure 1a). Conflicting phylogenetic signal in the data, however, is not sufficient to infer recombination (Huson and Bryant 2006). To test whether the NNet topology is due to recombination we used the pairwise homoplasy index (PHI) test (Bruen, et al. 2006), and the Benjamini-Hochberg correction for multiple testing (Benjamini and Hochberg 1995). Recombination signal (*q* ≪ 10^−6^) was detected among all included isolates (Figure 1a). SARS-CoV-1 and human SARS-CoV-2 monophyletic clades do not display net-like topology, and recombination signal of a sub-dataset including only these lineages was not significant (*q* = 0.14). SARS-like and bat-SL-CoVs, on the other hand, show a high degree of conflicting phylogenetic signals reflected by intricate networks (Figure 1a). SARS-like isolates are ancestors of SARS-CoV-1 isolates, bat-SL-CoVs stand as intermediate link between SARS-CoV-1 and SARS-CoV-2 isolates. Bat-SARS-CoV-2 RaTG13 (NCBI accession no. MN996532; GISAID id EPI_ISL_402131, isolated from bat in 2013 in China) and pangolin-SARS-CoV-2 isolates (EPI_ISL_410538-43 isolates denoted with “a”; and EPI_ISL_412860 and EPI_ISL_410721 isolates indicated as “b”, were isolated in 2017 and 2019 in China respectively, Figure 1a) are ancestors of SARS-CoV-2 circulating in humans. When scanning sub-sets of the alignment including only SARS-CoV-2 and its ancestors, recombination signal was highly significant (q < 1×10^−6^). A signal for recombination was also detected for SARS-CoV-2 with pangolin-SARS-CoV-2 ancestral isolates (q = 0.03), and when testing SARS-CoV-2 bat (EPI_ISL_402131) and pangolin isolates (q = 0.002). We next scanned the SARS-CoV-2 genomes for putative recombination breakpoints using RDP, GENECOV, MAXCHI, CHIMAERA, SISCAN, and 3SEQ, implemented in RDP4 (Martin, et al. 2015). This pipeline allows for the identification of recombinants as well as potential their parental isolates - if present in the sampled sequences. Sixty-eight potential recombination events distributed along the genome were found in bat-SL-CoVs, SARS-CoVs, and SARS-like CoVs isolates (Table S1). We found that human SARS-CoV-2 isolates were potentially either parental or recombinant sequences in recombination events with pangolin-SARS-CoV-2 isolated in 2019 and bat-SARS-CoV-2 genomes. Uncertainty in the estimation (the parental may be the recombinant sequences, or vice versa) suggests missing links in SARS-CoV-2 lineages, which is likely, given the sparse sampling of the wildlife coronavirus diversity, or low sensitivity of the recombination algorithms. Because the recombination event between CoVs of pangolin “b” and human origin involved the region of the S glycoprotein that binds to the cell receptor (Figure 1b-c and Table S1), exactly matching the ACE2 binding region where residues are shared between pangolin and human (Liu, et al. 2020), we conclude that the SARS-CoV-2 human lineage is the result of recombination between a strain belonging to the pangolin “b” lineage, potential minor parental, with a strain belonging to the bat lineage close to bat-SARS-CoV-2 RaTG13.

**Figure 1.**
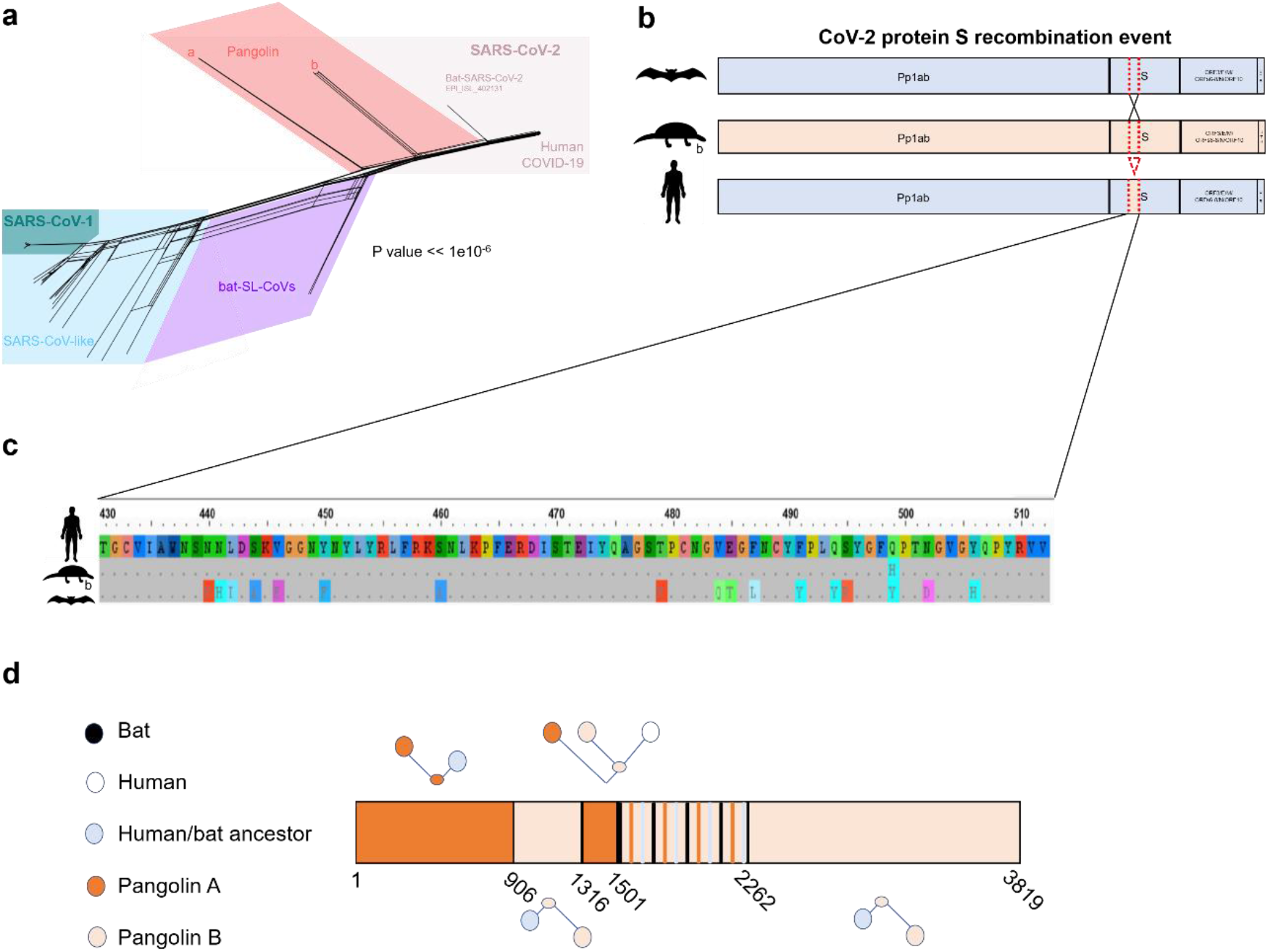
Recombination analysis across CoV genomes. (a) NNet graphs obtained using split-decomposition and based on the full genome sequence to detect recombination. P-value of PHI test of recombination is reported. Pangolin a taxa: EPI_ISL_410538-43 isolates isolated in 2017 in China; pangolin b taxa: EPI_ISL_412860 and EPI_ISL_410721 isolates isolated in 2019 in China. (b) Recombinant event involving human, bat and pangolin SARS-CoV-2 glycoprotein S. (c) Detail of the RBD recombinant fragment amino acid alignment. Dots represent identity with he human CoV-2. (d) Mosaic structure of SARS-CoV-2 glycoprotein S. The 4^th^ partition (1501-2262), as identified by GARD analysis, still presents some recombination signal (see table S4 for details).

We corroborated the potential recombination event by phylogenetic inference based on the recombinant and non-recombinant genome fragments from human, bat, and pangolin CoV-2 sequences, after assessing for the presence of phylogenetic signal (Table S2 and S3). Trees inferred using the recombinant region (part of the S glycoprotein) supported pangolin and CoV-2 ancestral relationship, while the segment derived by the major parent kept the CoV-2 clade clustering with the bat sequence (Figure S1).

We investigated further recombination patterns with a very sensitive genetic algorithm, GARD, based on a likelihood model selection procedure that searches multiple sequence alignments for evidence of recombination breakpoints (Kosakovsky Pond, et al. 2006). Extensive simulations have shown that such an in-depth analysis can detect recombination events with higher power and accuracy than tools based on phylogeny discordance among large genomic segments. Indeed, GARD detected a more complex mosaic-like structure of the S gene (Figure 1d), comprised of at least four distinct non-recombinant segments, in addition to a segment (S gene nucleotide position 1502-2262) with statistically significant evidence of further intra-segment recombination (Table S4). The mosaic is the result of multiple independent events involving different ancestral lineages during the evolutionary process that eventually gave rise to the current pandemic strain. The first two non-recombinant segments of the S gene (nt 1-906 and nt 907-1316) show a human/bat ancestor (that subsequently splits in Bat-SL-CoVs and SARS-CoV-2) sharing a common ancestor with Pangolin A and Pangolin B CoVs, respectively. The third segment (nt 1317-1501), which overlaps with the one identified by RDP4 analyses involving the RBD domain (Table S4), confirms its SARS-CoV-2/pangolin B CoVs common ancestry. The fourth segment (nt 1502-2262) appears to be itself a complex mosaic, although exact location of each putative breakpoint could not be calculated given the complexity of all possible patterns. The fifth segment (nt 1502-2262) shows, again, the human/bat ancestor sharing a common ancestor with Pangolin B CoVs. Overall, we hypothesize that the evolutionary past of the currently circulating pandemic strains included multiple recombination events between an ancestor of human SARS-CoV-2, or a closely related to bat-SARS-CoV-2, and an ancestor of pangolin-SARS-CoV-2 lineage b (but not necessarily in a pangolin host). Given that some regions in the SARS-CoV-2 genome show similarity to these clades (Figure S2), it is likely that one or more missing links among these strains have yet to be sampled in the animal reservoir.

### Prediction of recombination hot- and cold-spots in SARS-CoV-2

Recombination hotspots analysis was performed through a permutation test in RDP4, which randomly distributes the inferred breakpoints across the genome on the variable nucleotide positions between the three sequences involved in the event. From the permutations, density plots are drawn, and the distribution of observed breakpoints is compared to the permutations results. Permuting the detected breakpoints relative to variable positions takes into account the fact that breakpoints might be easier/more difficult to detect based on the local nucleotide diversity. Segments that had more observed breakpoints than the local maximum number of breakpoints of 99% of 1,000 permutations were considered hotspots (p=0.01); in the same fashion, segments that had fewer breakpoints than the local minimum of 99% of permutations were considered coldspots (Heath, et al. 2006). The algorithm uses a sliding window of a set size, which is moved across the genome one nucleotide at the time. The size of the sliding window may affect the sensitivity of the test, since smaller window sizes might detect tightly clustered hotspots, but might miss dispersed hotspots or coldspots. Thus, we used the windows size range 100-200 as suggested by the program authors. We plotted the locations of inferred recombination breakpoints and analyzed their distribution through permutations to predict hot- and coldspots of recombination across the coronavirus genomes (Figure 2).

**Figure 2.**
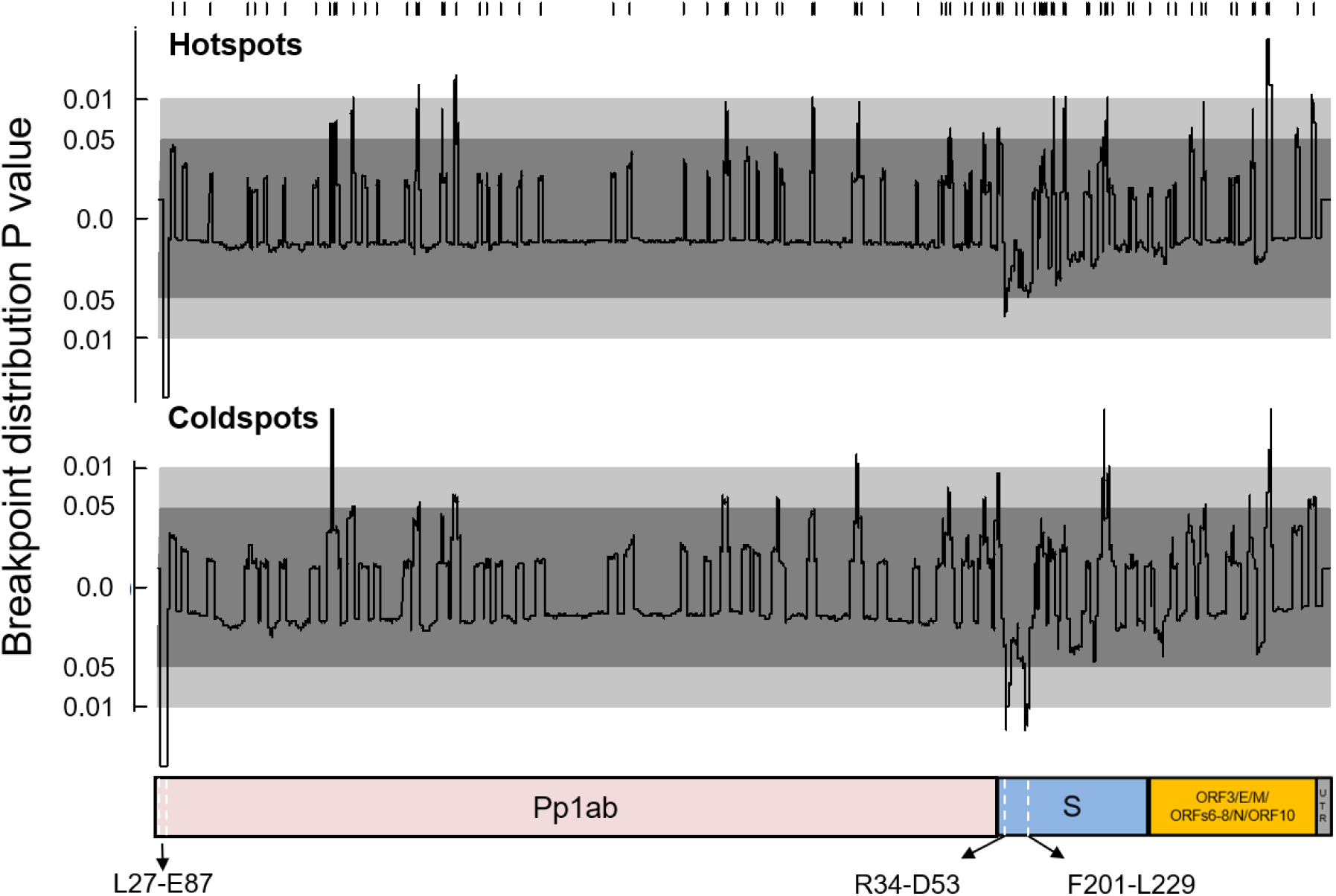
Recombination hot- and cold-spots prediction across CoV genomes. Hotspot (upper graph, sliding window size = 100) or coldspot (lower graph, sliding window size = 200) analysis. Detected recombination breakpoints are marked on the top of the panel. P-values ≤ 0.01 above the 0 mark indicate presence of hotspots (upper graph, “peaks”); in the bottom panel, p-values ≤ 0.01 below the 0 mark (“valleys”) indicate the presence of coldspots.

We identified nine potential recombination hotspots based on an analysis with sliding windows of size 100 bp. Seven of them were located in the polyprotein segment (Figure 2), which spans three quarters of the viral RNA; the remaining two involved protein E, *orf8*, and protein N. It has been reported that the *orf8* gene plays a role in regulating the viral replication in the host cell (Muth, et al. 2018), and was probably acquired by the whole lineage of SARS-CoV-1 from the SARS-related CoV from Greater Horseshoe bats (SARSr-Rf-BatCoVs) following a recombination event with a Chinese horseshoe bat coronavirus (SARSr-Rs-BatCoV) (Lau, et al. 2015). Three recombination coldspot regions were detected using a sliding window of size 200 (Figure 2). Despite the S glycoprotein being recombinant in nature, the N-terminal region holds two coldspots away from the recombination breakpoints: the first is located between residues R34 to D53, while the second between residues F201 to L229, (p ≤0.01). Another coldspot was found in the N-terminal region of Pp1ab. Coldspots might be possibly related to maintaining specific segments of the protein, or to the function of the viral transcription regulatory sequences (Kim, et al. 2020).

### SARS-CoV-2 spike (S) glycoprotein acquired unique features driven by recombination and purifying selection

We provide a full description of all residues unique to the S glycoprotein of SARS-CoV-2 isolates circulating among humans by comparison with SARS-CoV-2 from bat and two pangolin (lineage “a” isolates isolated from pangolin in 2017 in China; and “b” isolates isolated from pangolin in 2019 in China) (Table S5). The overall structure of the SARS-CoV-2 has significant structural homology to its SARS-CoV-1 counterpart (Wang, et al. 2020). In comparison with the latter available structure (Gui, et al. 2017), and a recently published study (Lan, et al. 2020), the SARS-CoV-2 S glycoprotein is composed of the S_1_ subunit, that binds to hACE2 (Li 2016), and the S_2_ subunit which plays a role in membrane fusion (Bosch, et al. 2003). In particular, S_1_ is composed of an N-terminal domain (NTD, residues 14-294 in SARS-CoV-1 and 14-303 in SARS-CoV-2) and three C-terminal domains (CTD1, CTD2, and CTD3) (Figure 3). The CTD1 (residues 320-516 for SARS-CoV-1 and residue 333-529 for SARS-CoV-2, see Methods) functions as the RBD, which allows SARS-CoV-2 to directly bind to the PD of the hACE2 (Li, et al. 2005). For simplicity, we show canonical hACE2 binding residues F486, Q493, S494 and N501, as reported by Andersen *et al* (Andersen, et al. 2020). The S_2_ subunit is located downstream of S_1_ after the S_1_/S_2_ cleavage site (residue 667 for SARS-CoV-1 and residue 684 for SARS-CoV-2). The majority of the residues that differentiate human SARS-CoV-2 S glycoprotein from bat or pangolin ones were found within the S_1_ subunit (Figure 3a, Table S6). While the S glycoproteins of MERS-CoV, HcoV-OC43, and HKU1 display the canonical (R/K)-(2X)n-(R/K)* (or RXXR*SA) motif for the S1/S2 cleavage (Li 2016; Gui, et al. 2017), the S glycoprotein of SARS-CoV-2 presents a solvent exposed furin-like cleavage sequence (PRRARS*V) in positions 681-684 (Andersen, et al. 2020), similar to the pattern found in a recently collected bat isolate (Zhou, Chen, et al. 2020) (Figure 3). The cleavage site within the S1/S2 junction of the S glycoprotein is the target for furin protease and essential for SARS-CoV-2 infection of humans (Hoffmann, et al. 2020). The high activity of the human furin, compared to bat, may enhance the kinetic of the infection, as the cleavage of S1 and S2 is a crucial step for virus entry into cells. For selection analyses, we retained only unique sequences (removing all other identical sequences) resulting in an alignment of 119 sequences. Because of the negative impact of recombination events on selection analyses (Schierup and Hein 2000; Posada and Crandall 2002; Shriner, et al. 2003), we used the additional partitioning identified by GARD for the selection analyses. On lineages that separate host clades (Figure 4), the S glycoprotein is under strong purifying selection (mean dN/dS = 0.0165), and 582 variable residues out of 838 found to be evolving with dN/dS < 1 (p-value ≤ 0.1) (Kosakovsky Pond and Frost 2005), including 106 residues out of 197 located in the RBD (Table S7). The recombination coldspots hosted a proportion of sites under purifying selection similar to the rest of the alignment. There was no evidence of episodic diversifying selection, either at the level of the entire alignment (dN/dS = 0.0 with weight 86.71%, dN/dS = 0.09 with weight 13.28%, p = 0.5) (Murrell, et al. 2015), or at individual sites (p < 0.1) (Murrell, et al. 2012), and no evidence was found of lineage specific episodic adaptation (p < 0.05). Focusing on internal branches for selection analyses removes the biasing effects of unresolved intra-host evolution or sequencing errors (Smith, et al. 2015). To identify sites which may be evolving adaptively in the SARS-CoV-2 clade, we tested for episodic diversifying selection only for internal branches in the human clade (Murrell, et al. 2012) and identified twenty-one such sites, eight in S_1_ (p < 0.1) (Figure 4, Table S6). Next, we investigated whether residues were evolving under the same selective pressure (Poon, et al. 2007). A total of five pairs of codons were found to coevolve together at posterior probability ≥ 0.9 (Figure S3); some of these residues are involved in epistatic interactions between residues in the RBD and external residues.

**Figure 3.**
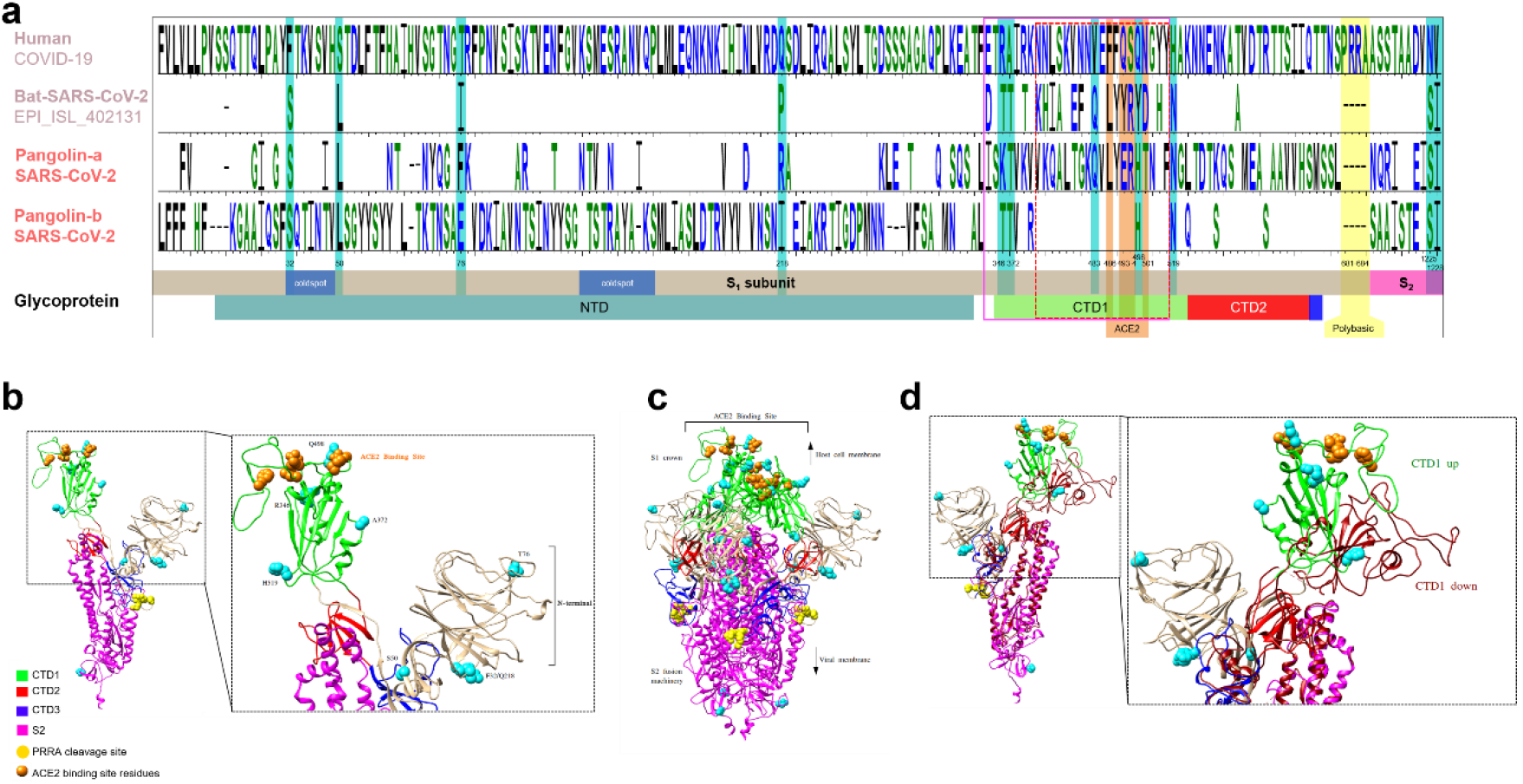
Signature residues in the S glycoprotein of SARS-CoV-2 and ribbon representations of the S protomer. (a) Signature residues that distinguish human SARS-COV-2 from closely related bat and pangolin (lineage ‘a’ isolates isolated in 2017 in China; ‘b’ isolates isolated in 2019 in China) isolates; residues are colored according their charge: hydrophilic (RKDENQ) residues in blue, neutral (SGHTAP) in green, and hydrophobic (YVMCLFIW) in black. Numbers correspond to the residue position in Wuhan-Hu-1 MN908947.3 isolate. In the box below we show a schematic representation of the spike glycoprotein is given: S_1_ and the S_2_ subunits of the S glycoprotein are indicated in grey and fuchsia, while CTD1 is shown green, CTD2 in red, CTD3 in blue. Blue boxes show coldspots. Fuchsia box around the residues between 333-516 are indicating the residues involved into rotation of RBD, red dotted box indicates residues 340-511 that SARS-CoV-2 obtained from the recombination event with pangolin lineage “b”. Residues unique to human SARS-CoV-2 are shaded in cyan, residues that form the hACE receptor binding are shaded in orange, and the acquired polybasic residues (PRRA) in the S_1_/S_2_ cleavage site in yellow. (b) The monomer of SARS-CoV-2 S glycoprotein and its three domains of the S_1_ subunit (CTD1, CTD2, and CTD3) and S_2_ subunit (pink) are shown in different colors: CTD1 in green, CTD2 in red, CTD3 in blue. The N-terminal domain was shown is colored tan. The residues unique to human SARS-CoV-2, hACE2 biding site residues, and S_1_/S_2_ cleavage sites were shown with cyan, orange and yellow spheres, respectively. (c) Homo-trimer structure of the modeled SARS-CoV-2. Residues, S_1_ (NTD, CTD1, CTD2, and CTD3) and the S_2_ subunits are colored corresponding to panel b. (d) The superposition of the modeled S protomer of SARS-CoV-2 with the S protomer of SARS-CoV (PDB 5WRG). The missing NTD domain for SARS-CoV was not shown whereas the NTD domain of SARS-CoV-2 is shown by tan color. Residues and domains are colored corresponding to panel b.

**Figure 4.**
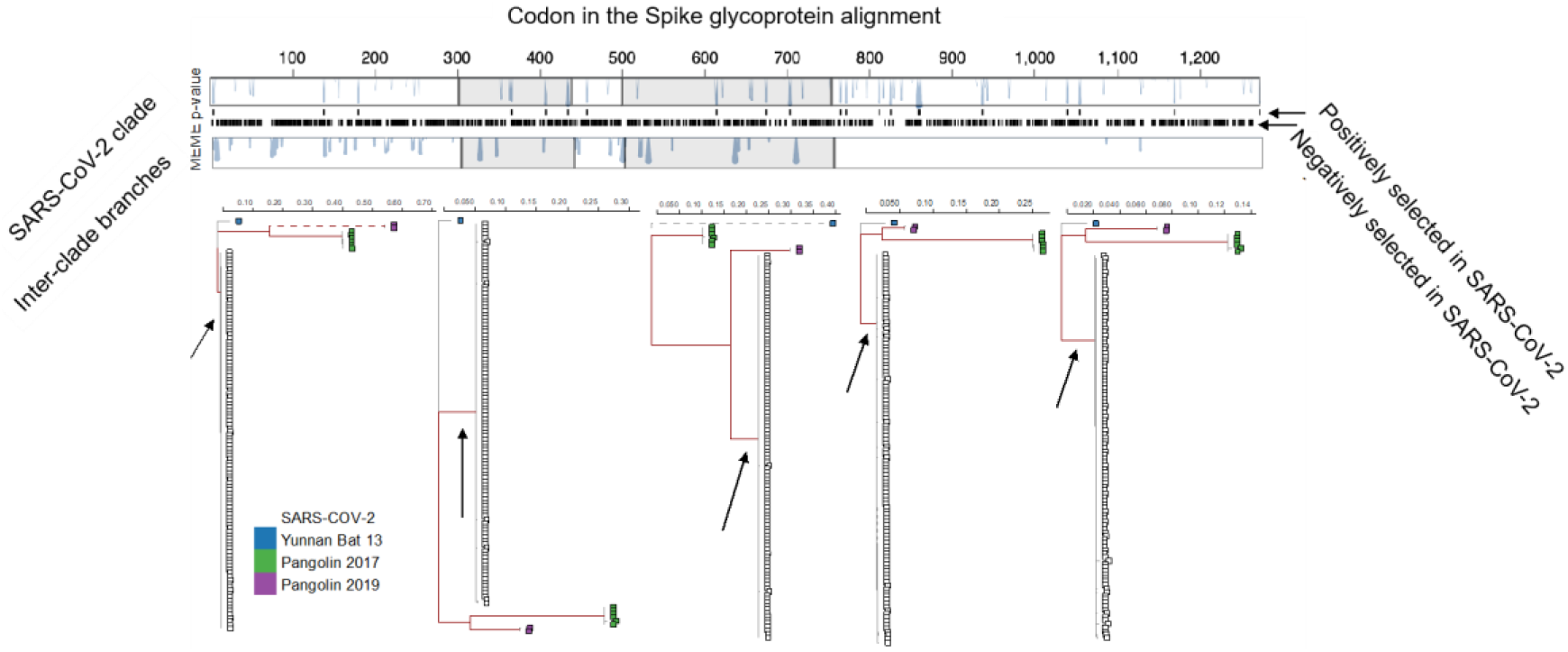
Selection analyses of Spike glycoprotein. Selection analyses were applied to a partitioned alignment of 119 CoV sequences, with partitions indicated by alternating white and grey rectangles. A maximum likelihood tree (rooted on the bat isolate for clarity) for each partition is shown, and the four inter-clade branches representing potential zoonotic events depicted in dark red. The branch ancestral to SARS-CoV-2 isolates has an arrow pointing to it. Branches longer that 0.25 subs/site (under the MG94 codon model in aBSREL analyses) are censored at 0.25 subs/site and shown in dashed lines. The impact of selective forces at individual sites is shown in two vertical bars at the top, where MEME p-values are shown either for the SARS-CoV-2 clade (top bar) or the inter-clade branches (bottom bar) as trail plots. Tick-marks between the bars correspond to the location of sites that were inferred to be subject to diversifying positive selection in the SARS-CoV-2 clade (there were none in the inter-clade branches), and those subject to pervasive purifying selection along the inter-clade branches.

### Polybasic furin-like cleavage and prefusion conformation correlate with “up” conformation

The RBD domain of SARS-CoV-2 S glycoprotein for the pre-fusion conformation rotates outward from center of the triangular head to an “up” position, while this conformation for SARS-CoV is “down”(Gui, et al. 2017; Wrapp, et al. 2020) (Figure 3d). The recognition step between SARS-CoV-2 RBD and hACE2 is a dynamic process in which the protein surface of both partners adapts one to the other, in line with cryo-EM evidences that shows how the receptor binding can open up RBD (Song, et al. 2018). In general, the RBD of each S monomer, prior to CoV infection, is buried in the inactive “down” conformation (prefusion state) and cannot bind to hACE2 due to steric clash (Wrapp, et al. 2020; Yan, et al. 2020). In the course of infection, one RBD monomer turned “up” to expose enough space to hACE2 (postfusion state), inducing further conformational open and loose for proteolysis (Gui, et al. 2017; Song, et al. 2018). Homology modeling of SARS-CoV-2 S glycoprotein showed a “up” conformation of one SARS-CoV-2 S monomer during the prefusion state, which was not showed for SARS-CoV-1.

We performed dynamic simulations (MD) to understand whether the genotype changes that we identified earlier played an important role in shaping the structure of the S glycoprotein of SARS-CoV-2, fundamental for viral binding to human cells. We carried out essential dynamics analysis on the MD trajectories, in order to separate the large collective protein movements connected to functional properties from the small and uninteresting motions (Amadei 1993). This analysis is based on the diagonalization of the covariance matrix built from the atomic fluctuations after the removal of the translational and rotational movement. It is usually applied on the c-alpha atoms only, since they describe the motion of the protein main chain. Interestingly, projection of the MD molecular trajectory along the first eigenvector (i.e. the direction that contains the majority of the protein total motion as shown by eigenvector values) is dominated by the up/down (opening/closing) motion of the RBD (see Figure 5a-b and Suppl. Movies 1-2). We identified threonine 333 (corresponding to T333 and T331 in bat and pangolin, respectively) as the hinge of this movement. Noticeably, we verified that this motion is conserved within the SARS-CoV-2 lineage, and therefore is observed in human, bat, and pangolin S glycoproteins (Figure 5f-g and h-i, respectively). It is also worth noting that residues 325-344 (mostly under negative/purifying selection) are nearly completely conserved between SARS-CoV-2, bat and pangolin CoVs (Figure 3a), in line with the key role of this protein region in guiding the up/down RBD movement.

**Figure 5.**
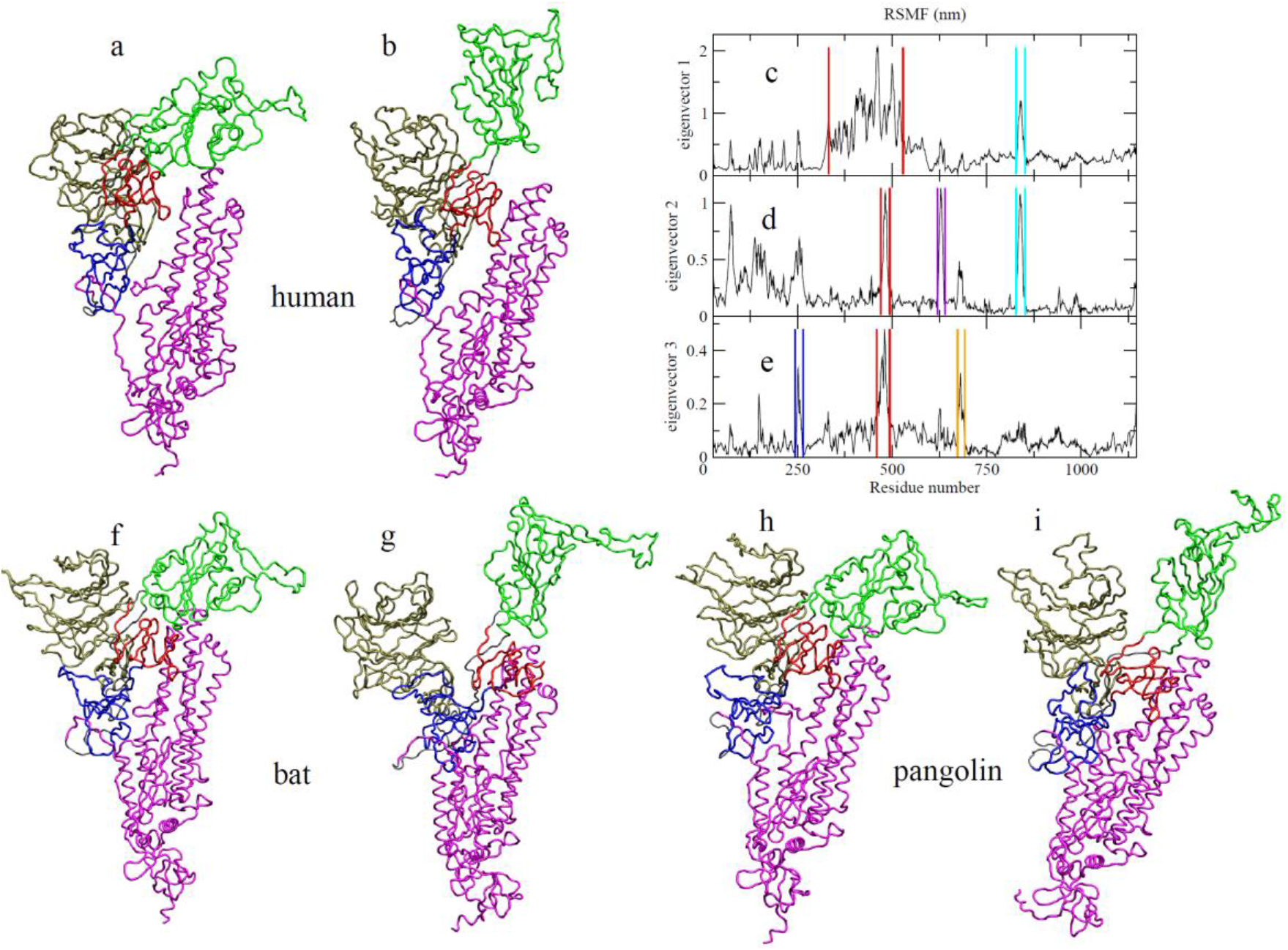
Essential Dynamics analysis of SARS-CoV-2 in comparison with CoV from bat and pangolin. (a-b) Projection of the SARS-CoV-2 S glycoprotein MD trajectory along Essential Dynamics (ED) eigenvector 1 (see also Supplementary movies 1-2). The two extreme conformations are shown in (a) and (b). Domain colors as in panel Figure 3b. The opening/closing mechanism of the SARS-CoV-2 S glycoprotein monomer by means of the rotation of RBD is evident. (c) Per-residue Root Mean Square Fluctuations (RMSF) in nm of the SARS-CoV-2 S glycoprotein filtered trajectory along ED eigenvectors 1. The region with higher fluctuations is the CTD1, highlighted by red lines, demonstrating that its rotation in the opening/closing mechanism is the most correlated motion in the protein. The peak in fluctuations involving residues 828-852, highlighted by cyan lines, demonstrates that when the RBD is in the up conformation, the fusion peptide visits a more solvent exposed conformation (see Figure S4 and Supplementary movie 3); (d) as in (c) but along eigenvector 2. The peak of fluctuations highlighted by red, purple and cyan lines are indicative of a direct structural communication between the hACE2 binding region, residue 620-640 in CTD3 and the fusion peptide (see Figure S5 and Supplementary movie 4); (e) as in (c) but along eigenvector 3. The peak of fluctuations highlighted by blue, red and orange lines are indicative of long range communications between residue 244-265 in NTD, the hACE2 binding residues and the furin-like cleavage site (see Figure S6 and Suppl Movie 5).

The conformational flexibility of the S protein during the sampled 100 ns trajectory was assessed by means of root mean square fluctuations (RMSF), calculated after superimposing each individual structure of a trajectory onto the initial structure by means of least-squares fitting, to remove rotational and translational motions. Projection of the MD molecular trajectory along the first three eigenvectors (that cumulatively contain 90% of the total protein motion) allowed to highlight co-variant movements (simultaneously changing) among long distance S glycoprotein regions, functional to the protein rearrangement during the pre-fusion step (Figure 5c-e). In detail, projection along eigenvector 1 (Figure 5c) shows a correlated movement between RBD and the region 828-852 (highlighted by cyan lines in Figure 5c), encompassing the C-terminal portion of the fusion peptide, defined by residues 816-833. The up conformation of RBD, in particular, corresponds to a more solvent exposed conformation of the fusion peptide, that is functional to the fusion of the viral and the host cell membranes (Figure S4 and Suppl. Movie 3). Projection along eigenvector 2 (Figure 5d) shows a correlated movement between a more restricted portion of RBD (residues 470-494, highlighted by red lines in Figure 5d), and again the previously defined region 828-852. A third peak in RMSF along this eigenvector is observed in the region 620-640, in CTD3. It is significant that the indicated portion of RBD includes three out of the four hACE2 binding residues, i.e. F486, Q493 and S494, demonstrating a direct structural communication between the ACE2 binding region and the fusion peptide (see Figure S5 and Suppl. Movie 4). Projection along eigenvector 3 (Figure 5e), finally, showed a correlated movement between residues 244-265 in the NTD region (blue lines in Figure 5e), residues 460-494 in CTD1 (red lines in Figure 5e), and residues 673-692 (orange lines in Figure 5e), which encompass the furin-like cleavage site typical of SARS-CoV-2. Also in this case, the up conformation corresponds to a more solvent exposed surface of the furin-like cleavage site. A peak of fluctuation for the furin-like cleavage site is observed also along eigenvector 2 (Figure 5d), even though of minor amplitude as compared to the highlighted three co-variant regions (i.e. hACE2 binding residues, fusion peptide and res 620-640 in CTD3) along eigenvector1. The long-range effects between the hACE2 binding residues and the furin-like cleavage site (see Figure S6 and Suppl Movie 5), might therefore be functional to the novel acquired capabilities of this virus. This motion, paired with the open/up conformation of SARS-CoV-2 glycoprotein that exposes the whole interaction interface of RBD and the fusion peptide to hACE2 and host cell membrane, might explain why SARS-CoV-2 has been shown greater infectivity and spread than previous CoVs.

## Discussion

Recombination is a hallmark of CoV evolution (Graham and Baric 2010). Although our results clearly point to several recombination events occurred between ancestral pangolin- and bat-SARS-CoV-2 lineages, the limited sampling of these animal hosts makes difficult to draw definitive conclusions about its origin. According to our results, one of these events led to the current configuration of SARS-CoV-2 ACE2 binding residues, including recently described residues forming hydrogen bonds with the host cell receptor (Yan, et al. 2020).

Current algorithms cannot fully distinguish recombinant sequences from the parental ones, at least for some recombination events. Yet, the lack of positive diversifying selection on the branch, in the S glycoprotein phylogeny, that leads to the recent human CoV-2 lineage, and the strong purifying negative selection exerted on the majority of the S glycoprotein codons, support our hypothesis that recombination constitutes the main drive to escape a local fitness maximum (Weinreich and Chao 2005), successfully expanding host tropism. It is likely that the ancestral lineages that actually recombined is part of a yet unsampled population. A more extensive sampling of wildlife CoVs would help in clarifying the events that led to the jump to humans. Our analyses were based on sequences obtained from strains circulating relatively early in the epidemic; future work including later strains’ genomes might discover marks of further adaptation to the new host, as new mutations emerge and get fixed in the human population.

Despite the frequency of recombination events involving the S glycoprotein, the N-terminal region appears to host two recombination coldspots. While mutation/recombination events in the glycoprotein RBD could result in expanded viral tropism, recombination coldspots and negative/diversifying selective pressure limit variability, thus maintaining essential structure and function. It must be noted that the high diversity between CoVs in the glycoprotein region, as compared to the rest of the alignment, might have an impact on this analysis, thus *in vitro* studies might be needed to verify these findings. Yet, while RBD is definitively a good target for therapeutic development, regions in coldspots or those undergoing negative/diversifying selective pressure may be helpful for successful vaccine design.

SARS-CoV-2 S glycoprotein gained specific features that have facilitated the successful spread of the virus as compared to its predecessors.

SARS-CoV-2 S glycoprotein presents two notable changes: a favorable furin-like cleavage site (Andersen, et al. 2020), and RBD up in pre-fusion mode (Wrapp, et al. 2020). Furin is ubiquitously expressed at different levels in all tissues (Schalken, et al. 1987; Hatsuzawa, et al. 1990), but kinetics of furin-like enzyme activity are different among species – e.g., human lung cells display quicker kinetic than bat (Pteropus *alecto*) (El Najjar, et al. 2015) – and may not fully explain SARS-CoV-2 boosted infectivity. Thus, the assumption that the insertion of a furin-cleavage site allowed a bat CoV to gain the ability to infect humans facilitating the host specie jump once the conditions were favorable cannot be excluded. On the contrary S glycoprotein of previously circulating SARS-CoV isolates do not have a favorable furin-like cleavage site (Andersen, et al. 2020) and remains un-cleaved after biosynthesis (Belouzard, et al. 2009). Furin has the potential to cleave specifically viral envelope glycoproteins, to enhance viral fusion with host cell membranes (Izaguirre 2019), thereby any variation/mutational changes in amino acid composition at cleavage sites may impact tissue and cell tropism, host range, and pathogenesis (Klenk and Garten 1994), as reported for other respiratory viruses (Klenk and Garten 1994; Peng, et al. 2017; Izaguirre 2019). MD simulations correlated more solvent-exposed surface in furin-like cleavage site and fusion peptide the S protein up conformation, identifying a conserved region that acts as a hinge in the opening/closing mechanism of all the SARS-CoV-2 lineage. Development of inhibitors of such motion may reduce substantially pre-fusion conformation and therefore the infectivity of SARS-CoV-2. Finally, the presence of coevolving sites might derive from host tropism; recent *in silico* and *in vitro* studies have already found evidence of epistatic effects in coronavirus genes (Donaldson, et al. 2007; Stobart, et al. 2012; Forni, et al. 2016). These connections might be linked to host adaptation (Bhatt, et al. 2013; Longdon, et al. 2014; Bedhomme, et al. 2015). Further *in vitro* research is needed to verify these findings and investigate the driving biological mechanisms behind it.

In summary, our study provided novel molecular insights on SARS-CoV-2 glycoprotein features and placed SARS-CoV-2 lineage in a new context within the evolutionary trajectory that has led to the current pandemic. Extensive recombination of isolates circulating in bat, pangolin and/or potentially other unsampled hosts such as civets or pigs, has been a crucial factor facilitating the emergence of a highly infectious virus, capable both to elude the immune system (Yuan, et al. 2020) and infect target cells more efficiently (Andersen, et al. 2020). The landscape of CoVs diversity still needs to be exhaustively explored, and future emergence of novel recombinant CoVs, and other viruses as well, should be an urgent concern. Strengthening surveillance of viral populations and improving our understanding of their dynamics will help the prevention of new zoonotic outbreaks and spread of emerging pathogens.

## Materials and Methods

### Genetic data

Publicly available genomic CoVs sequences were obtained from GenBank and GISAID (Table S5 and S7). To gather closely related CoVs BLASTn was performed using the ViPR database and keeping the first 100 hits. Genome sequences were aligned using MAFFT (Katoh and Standley 2016) and refined manually. Alignments for the glycoprotein were extracted from the genome alignment.

### Recombination breakpoint and hotspot analyses

The presence of conflicting phylogenetic signals (i.e. distinct tree topologies compatible with the same set of aligned sequences) in the full genome data set was investigated, first, by inferring networks using neighbor-net (Bryant and Moulton 2004) implemented in SplitsTree4 (Huson and Bryant 2006). Significant presence of recombination signal was then assessed with the pairwise homoplasy index (Phi) test (Bruen, et al. 2006). Identification of putative recombinant events and associated breakpoints was performed using the RDP(Martin and Rybicki 2000), GENECOV (Padidam, et al. 1999), MAXCHI (Smith 1992), CHIMAERA (Posada and Crandall 2001), SISCAN (Gibbs, et al. 2000) and 3SEQ (Boni, et al. 2007) algorithms implemented in RDP4 (Martin, et al. 2015) software (http://web.cbio.uct.ac.za/~darren/rdp.html), v.4.97. Statistical evidence of recombination was indicated by *p*-values < 0.05, after Bonferroni correction for multiple comparisons. A putative recombination events were considered as such if supported by at least five of the six algorithms used. Full table with recombination results id given in Table S1. BOOTSCAN (Salminen, et al. 1995) plots were generated using RDP4. Phylogenetic signal for the subset alignment including bat, human and pangolin CoV-2 sequences was l was confirmed with TREE-PUZZLE(Schmidt, et al. 2002), and weighted SH and AU tests (Shimodaira 2002) implemented in PAUP* v.4.04a build 167 (Hancock 2014). Neighbor joining, maximum likelihood and Bayesian trees were calculated respectively with Q-Tree v.2.0 (Nguyen, et al. 2015) and MrBayes (Ronquist, et al. 2012); best models for the alignments were chosen using Model Finder (Kalyaanamoorthy, et al. 2017).

Recombination hotspots analysis was also performed using RDP4 through a permutation test, which randomly distributes the observed breakpoints across the genome on the variable nucleotide positions between the three sequences involved in the event. From the permutations, density plots are drawn, and the distribution of observed breakpoints is compared to the permutations results. Two iterations were performed, with sliding windows of size 100 and 200, moved one nucleotide at the time, with different sensitivity to hotspots and coldspots. Segments that had more observed breakpoints than the local maximum number of breakpoints of 99% of 1,000 permutations were considered hotspots (p=0.01); in the same fashion, segments that had less breakpoints than the local minimum of 99% of permutations were considered coldspots (Heath, et al. 2006).

To identify the recombinant structure used to correct selection analyses, we used a genetic algorithm, GARD (Kosakovsky Pond, et al. 2006), on the subset of sequences including all bat and pangolin isolates, as well as five randomly chosen SARS-CoV-2 sequences; the reason for this subsampling is that for a fixed number of sites, GARD loses power when there are many nearly related sequences.

### Signature pattern and selection analyses

To identify new features of CoV-2, we used a higher number of isolates permit to explore a larger evolutionary landscape. VESPA algorithm (Korber and Myers 1992) was used with a dataset comprising the 728 genomes of SARS-CoV-2, and SARS-CoV-2 isolates from the closely related bat (EPI_ISL_402131, 2013, Accession MN996532.1) and pangolin lineage “a” (EPI_ISL_410538-43 isolates isolated in 2017 in China) and lineage “b” (EPI_ISL_412860 and EPI_ISL_410721 isolates isolated in 2019 in China) (Table S8). Positions are numbered with SARS-CoV-2 isolate Wuhan-Hu-1 (Accession MN908947.3) as reference (Table S9).

For selection analyses, we used GARD results to partition the 119 unique sequences subset. We inferred ML trees separately for each segment using *raxml-ng* and the GTR+G+I model with 5 random parsimony starting trees. In these trees, we further identified three branches that included host-switching events and the branch separating 2017 and 2019 pangolin isolates (see Figure 4).

We used BUSTED (Murrell, et al. 2015) to assess the presence of selection on the gene S partitions. We ran FEL (Kosakovsky Pond and Frost 2005) and MEME (Murrell, et al. 2012) methods on this partitioned alignment to look for pervasive (FEL) or episodic (MEME) diversifying positive selection affecting the four inter-clade branches, and, in a separate analysis, affecting only SARS-CoV-2 clade. To look for lineage specific selection on inter-clade branches we used the aBSREL (Smith, et al. 2015) method separately on each partition (since it cannot be applied to multi-partition data). Finally, to look for co-evolution between sites, we applied BGM (Poon, et al. 2007) to each partition separately, focusing only on internal branches and sites that accumulated at least two substitutions on internal branches. All selection analyses were carried out in HyPhy v2.5.14 (Pond, et al. 2005).

### Structure homology modeling

As a first step, the amino acid sequences of SARS-CoV-2 Wuhan-Hu-1 (Accession QHD43416) and EPI_ISL_402131 (bat, 2013, Accession MN996532.1) were submitted to Swiss-model (Waterhouse, et al. 2018). The best models showing high values of Global Model Quality Estimation (GMQE) were selected and used as templates. Amino acid alignments of targets and templates were performed using EMBOSS Needle (Madeira, et al. 2019). The 3D homology homo-trimer models were created with the Modeller v9. 23 (Eswar, et al. 2006; Webb and Sali 2014) automodel class using the alignment as a guide. Then Discreet Optimised Protein Energy Score (DOPE) based model selection (Shen and Sali 2006) and refinements were conducted using Modeller v9.23 scripts. The final structure model validation was conducted with ProSA (Sippl 1993; Wiederstein and Sippl 2007) and QMEAN servers (Benkert, et al. 2009; Benkert, et al. 2011). Visualization of the atomic models, including figures and movies, is made with Chimera v1.12 (Pettersen, et al. 2004) and VMD v1.9.2 (Humphrey, et al. 1996).

### Molecular Dynamics simulation

The structure of SARS-CoV-2, bat and pangolin, obtained for homology modeling, were used as starting structures for Molecular Dynamics (MD) simulations. In detail, topology file was generated with the pdb2gmx Gromacs tool (Páll 2015), using the amber99sb forcefield (Lindahl, et al. 2017)..The proteins were embedded in a triclinic box, extending up to 15 Å from the solute, and immersed in TIP3P water molecules (Jorgensen 1983). Counter ions were added to neutralize the overall charge with the genion gromacs tool. After energy minimizations, the systems were slowly relaxed for 5 ns by applying positional restraints of 1000 kJ mol^−1^ nm^−2^ to the protein atoms. Then unrestrained MD simulations were carried out for a length of 100 ns for each system, with a time step of 2 fs, using GROMACS 2018.3 simulation package (Páll 2015) (supercomputer Galileo, CINECA, Bologna, Italy). V-rescale temperature coupling was employed to keep the temperature constant at 300 K (Bussi 2007). The Particle-Mesh Ewald method was used for the treatment of the long-range electrostatic interactions (Darden 1993). The first 5 ns portion of each trajectory was excluded from the analysis.

We carried out the Essential Dynamics analysis to identify the main 3N directions along which the majority of the protein motion is defined (Amadei 1993). This analysis is based on the diagonalization of the covariance matrix built from the atomic fluctuations after the removal of the translational and rotational movement and we applied it on the c-alpha atoms of each monomer (residue 27-1146 in our model of SARS-CoV-2), after concatenating their trajectories (255 ns total sampling).

## Acknowledgements

MS is supported in part by the Stephany *W*. Holloway University Chair in AIDS Research. This work was supported by the Tunisian Ministry of Higher Education and Scientific Research, the ‘Departments of Excellence-2018’ Program (Dipartimenti di Eccellenza) of the Italian Ministry of Education, University and Research, DIBAF-Department of University of Tuscia, Project ‘Landscape 4.0–food, wellbeing and environment’. We thank all those who have contributed SARS-CoV-2 genome sequences to the GISAID database (https://www.gisaid.org/). We acknowledge Cineca and ELIXIR-IIB for computing resources We would like to thank Dr. Darren Martin for his advice on the recombination hotspot analysis.

