## Supplementary figures and images for "Recombination and purifying selection preserves covariant movements of mosaic SARS-CoV-2 protein S"

### figure s1

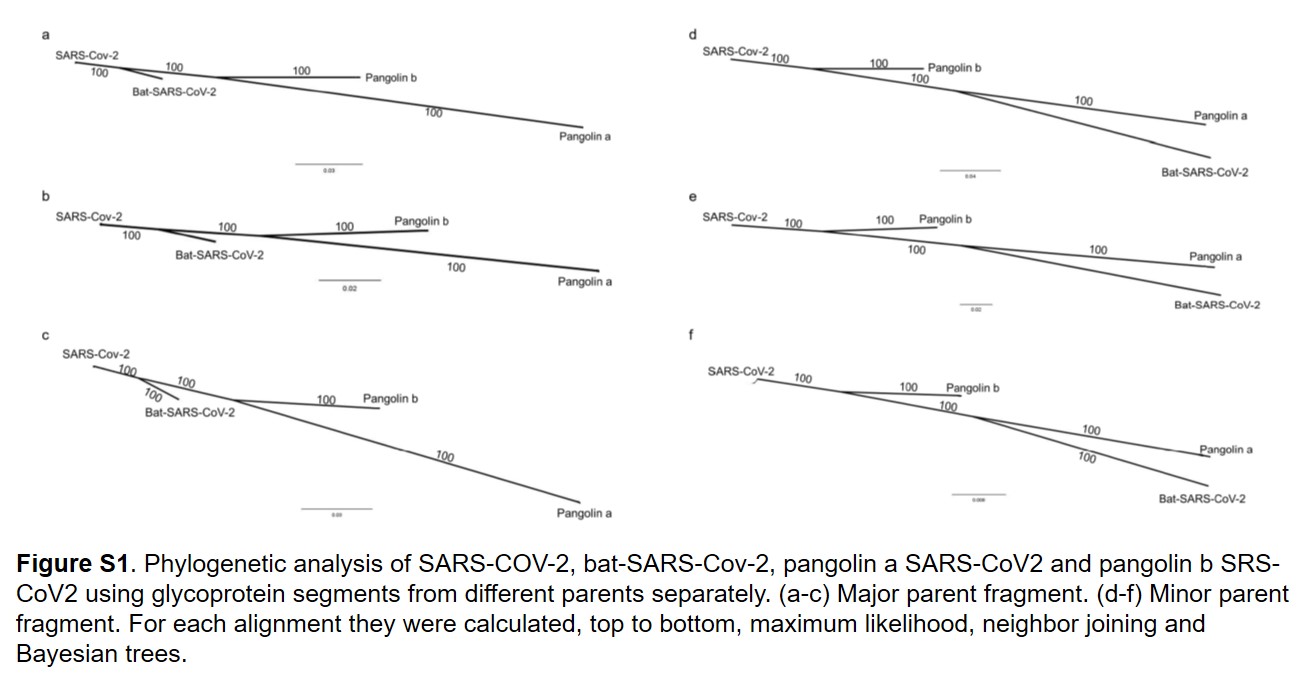

### figure s2

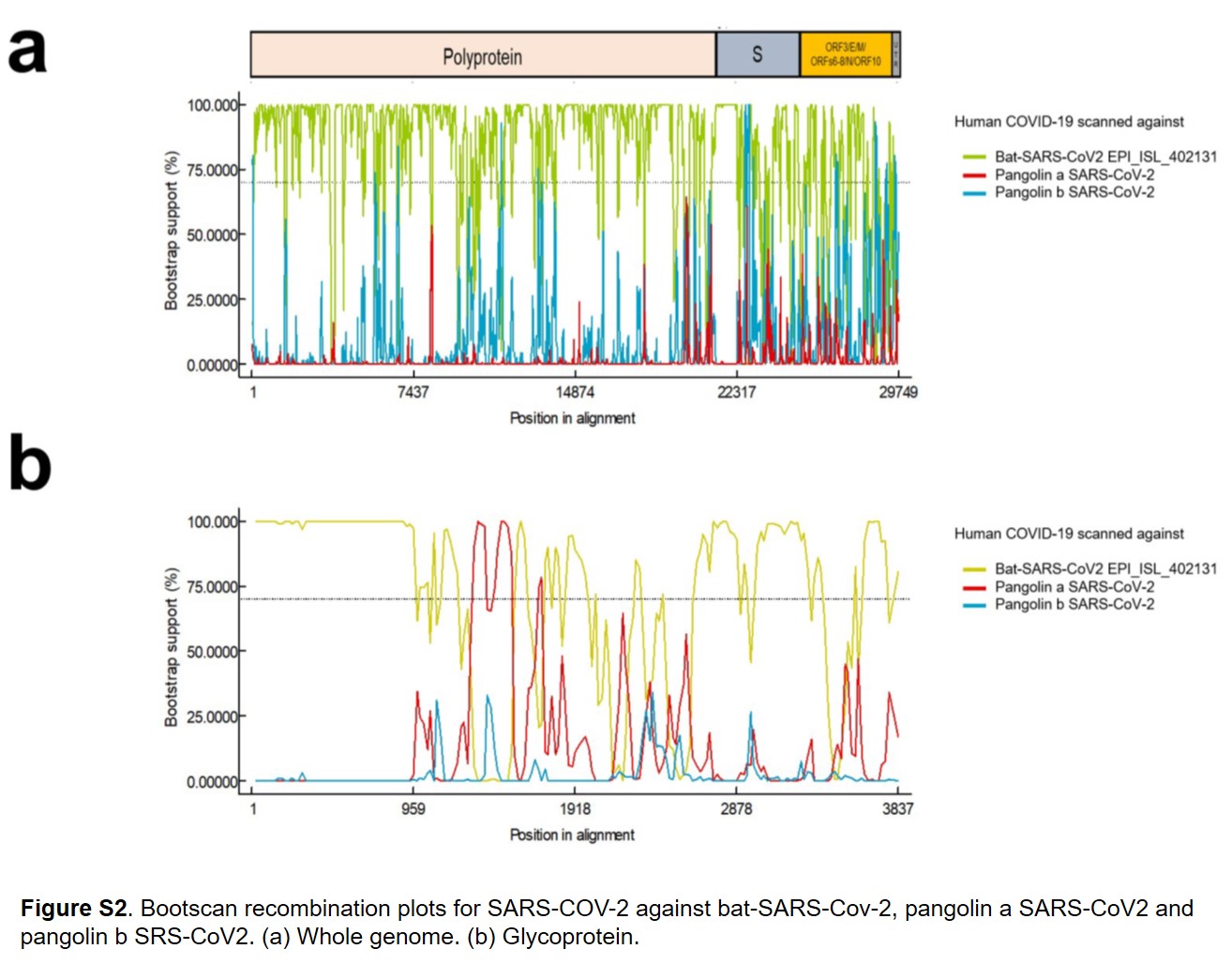

### figure s3

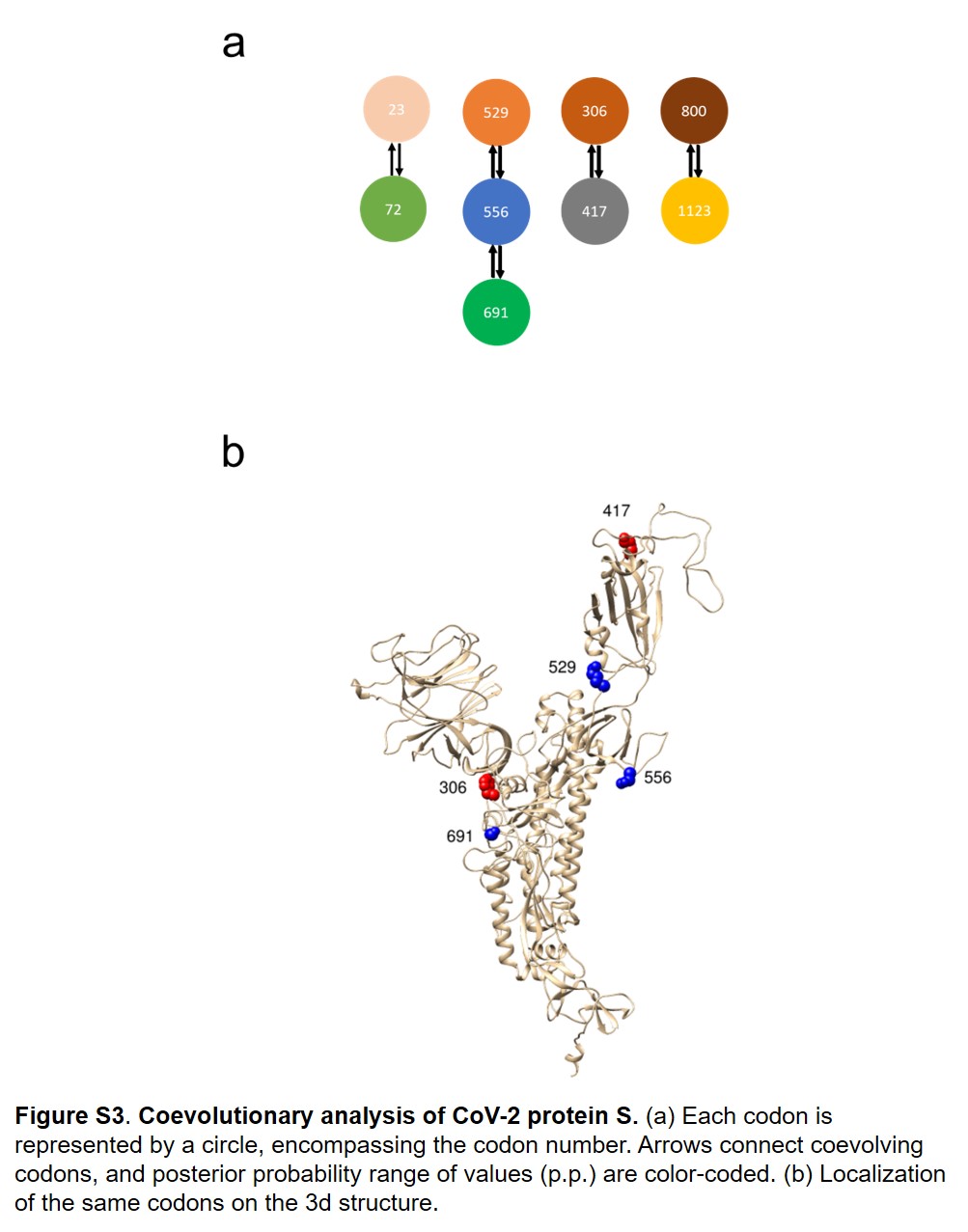

### figure s4

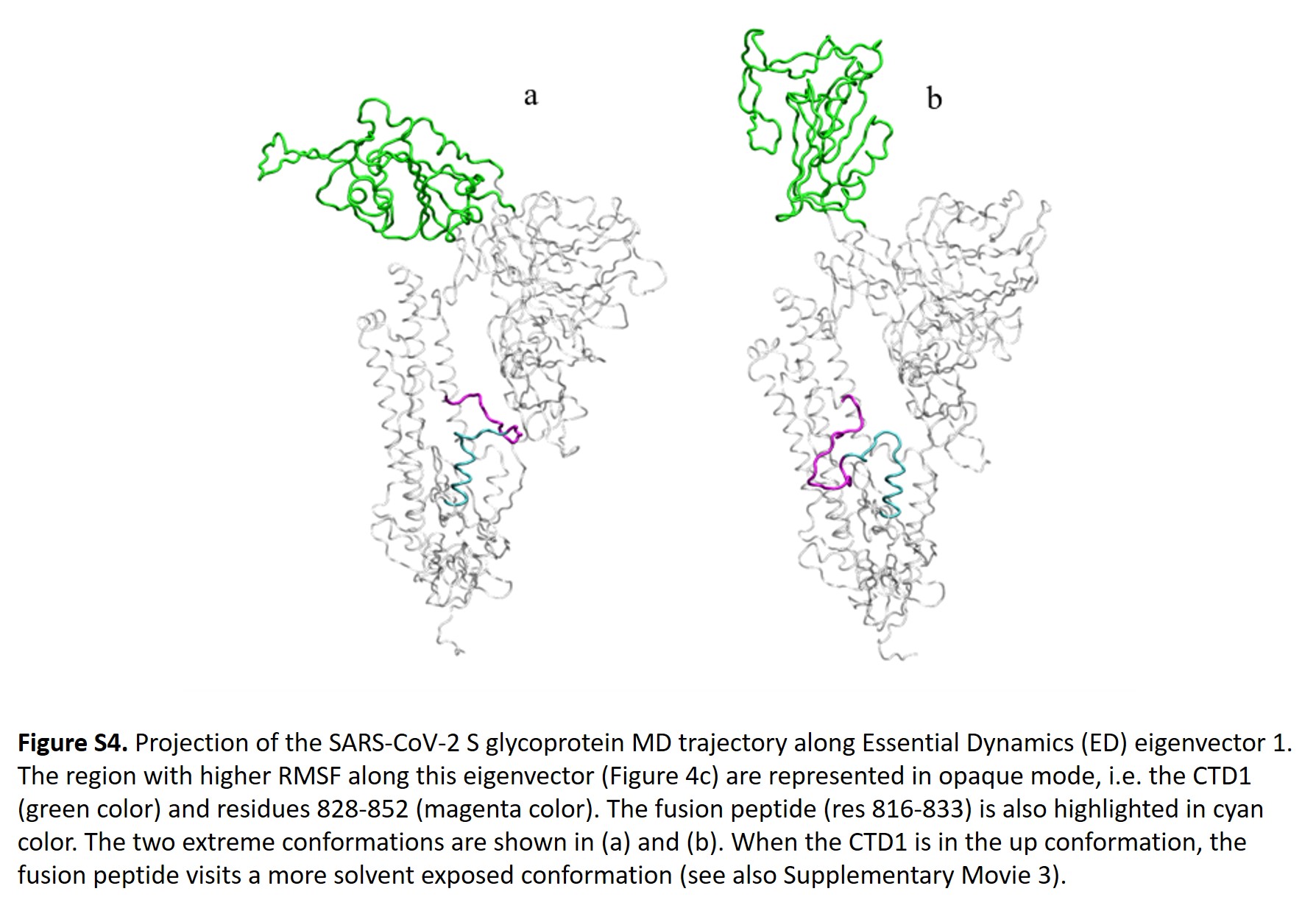

### figure s5

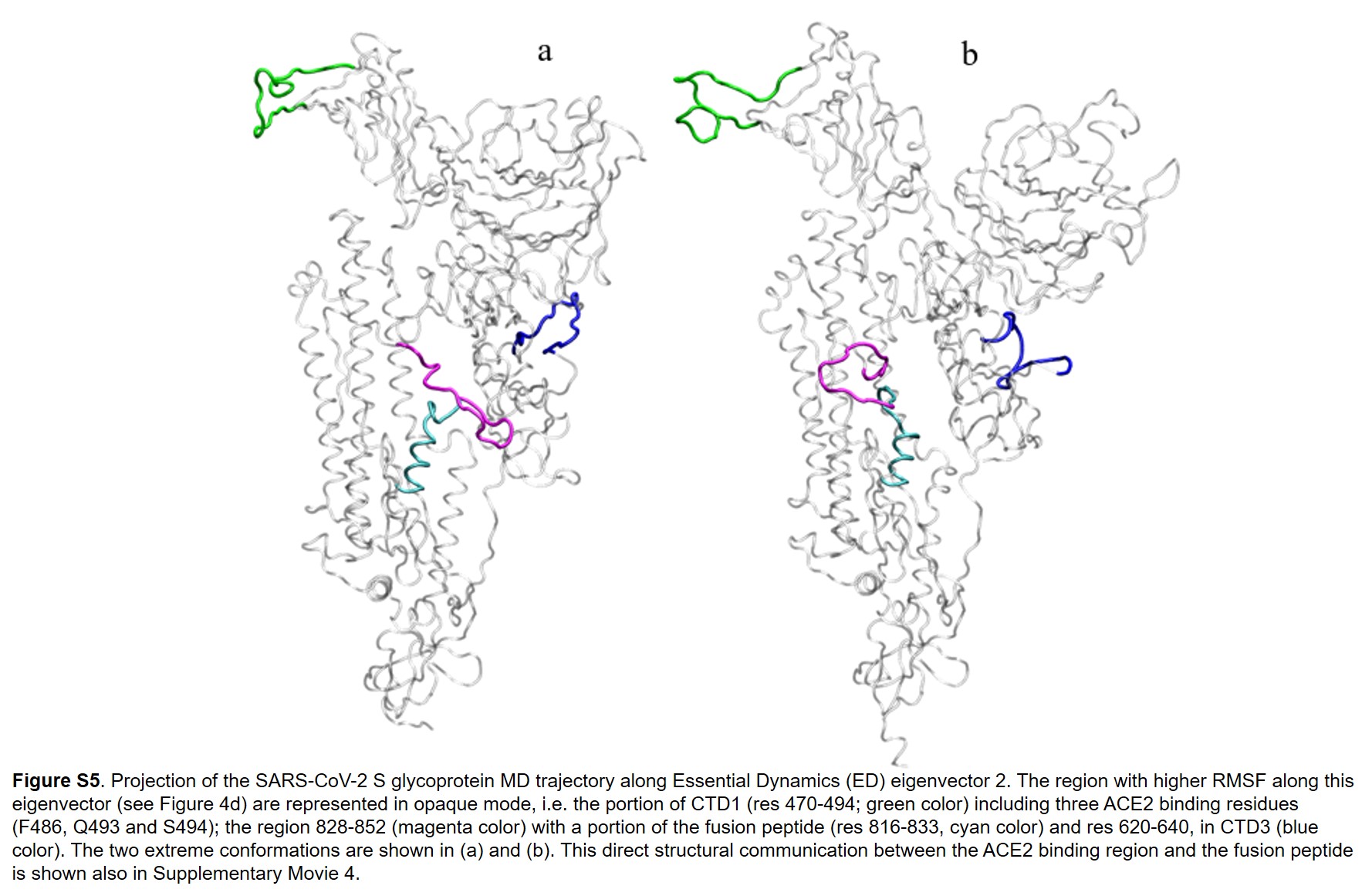

### figure s6

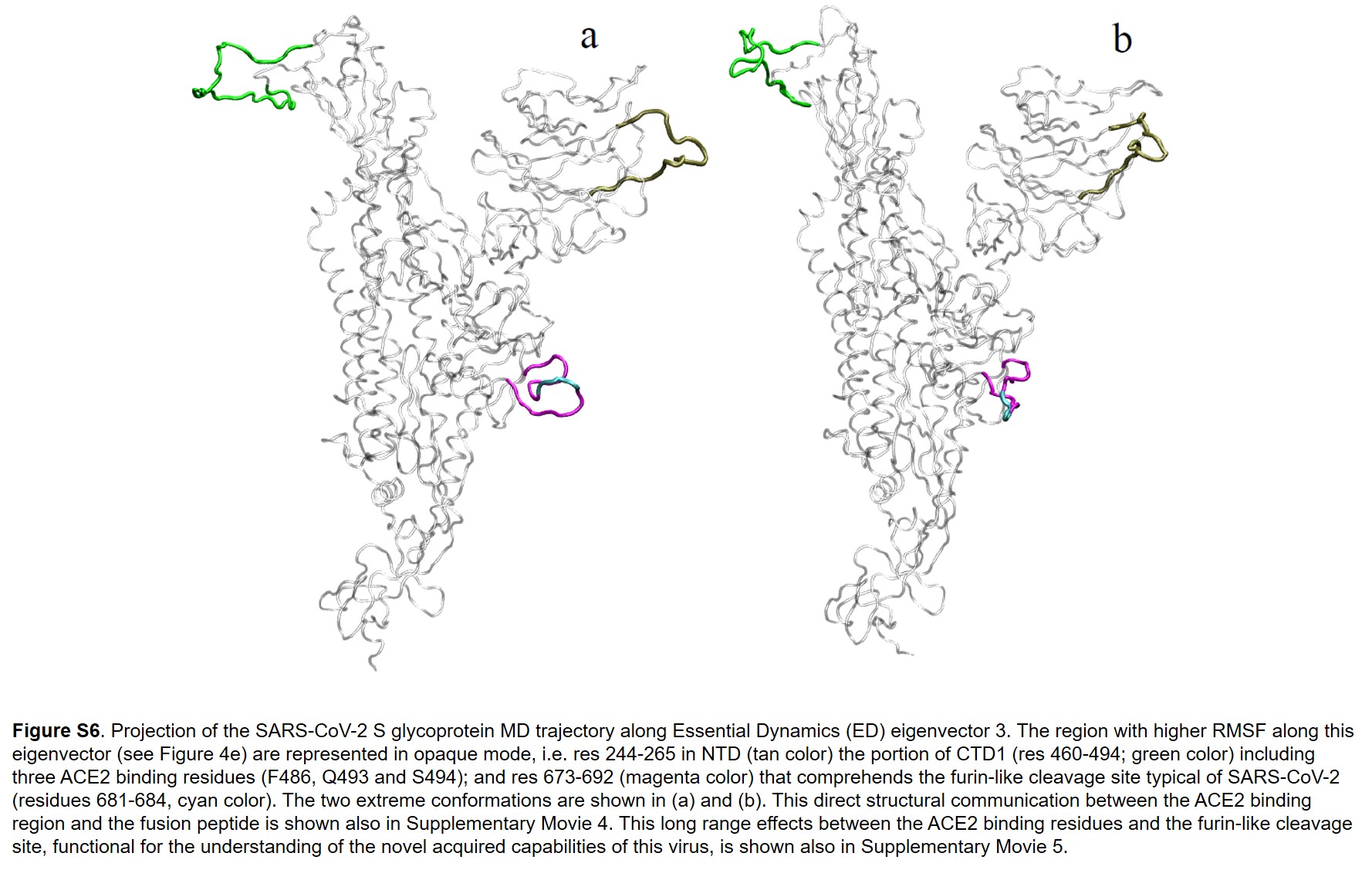

### movie 1

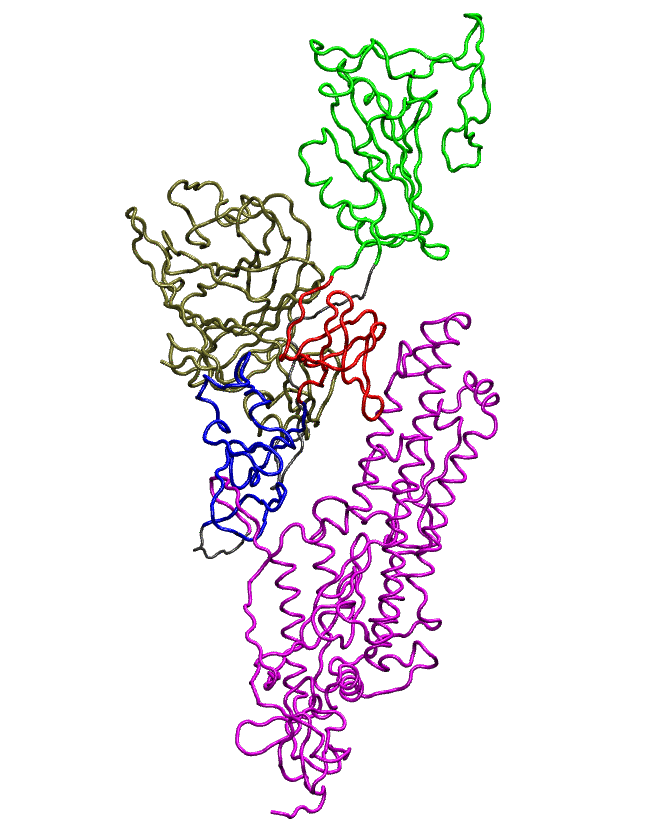

### movie 2

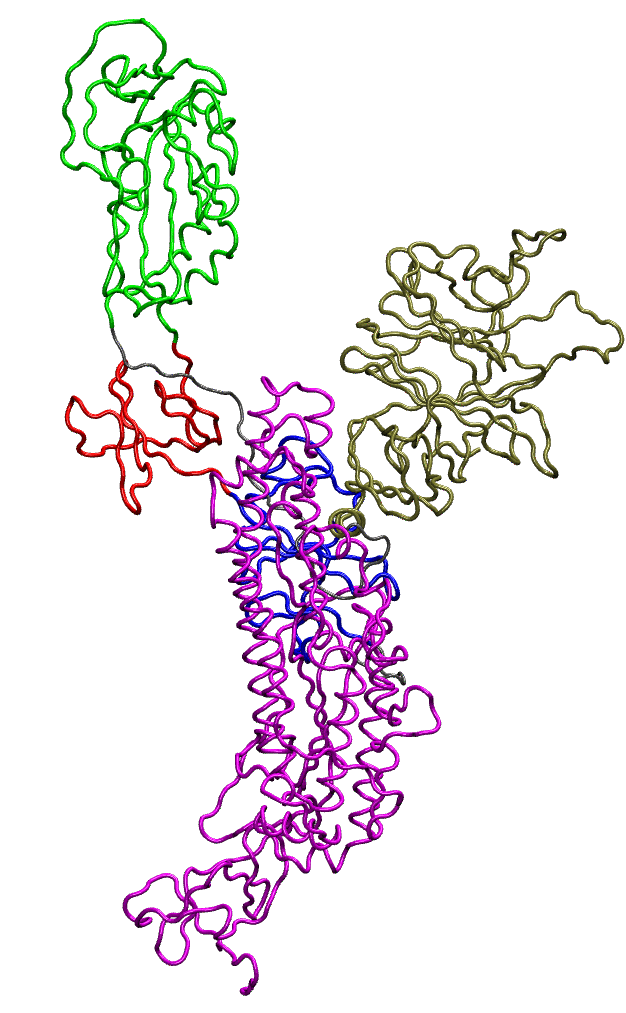

### movie 3

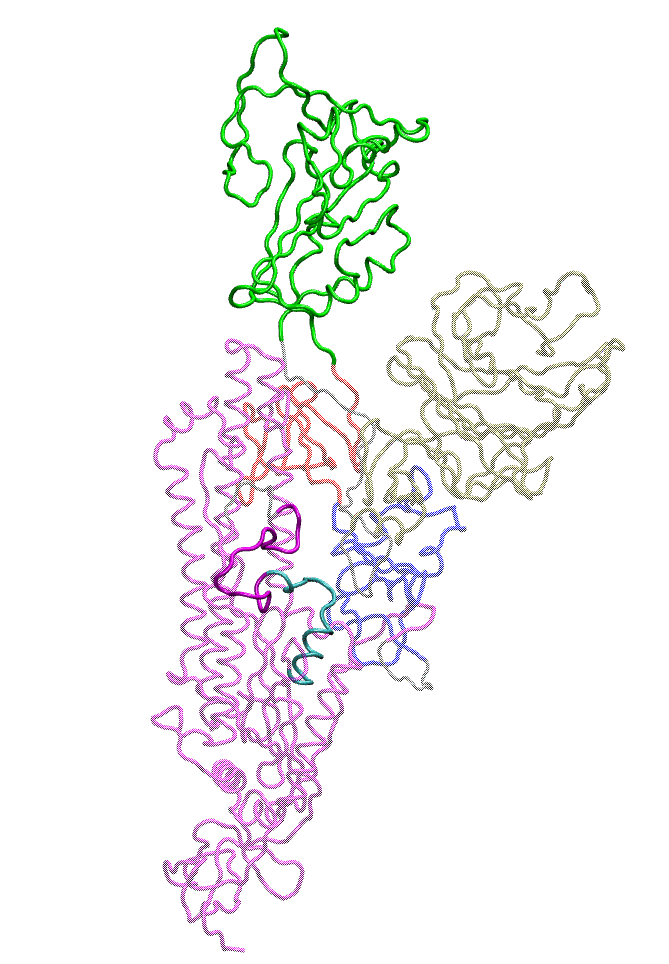

### movie 4

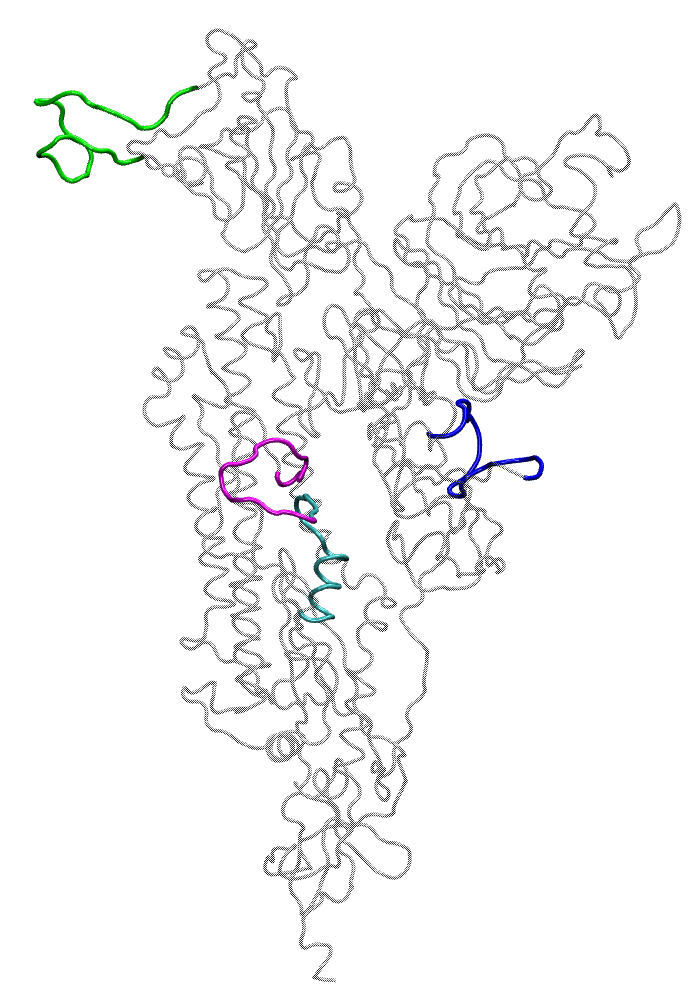

### movie 5

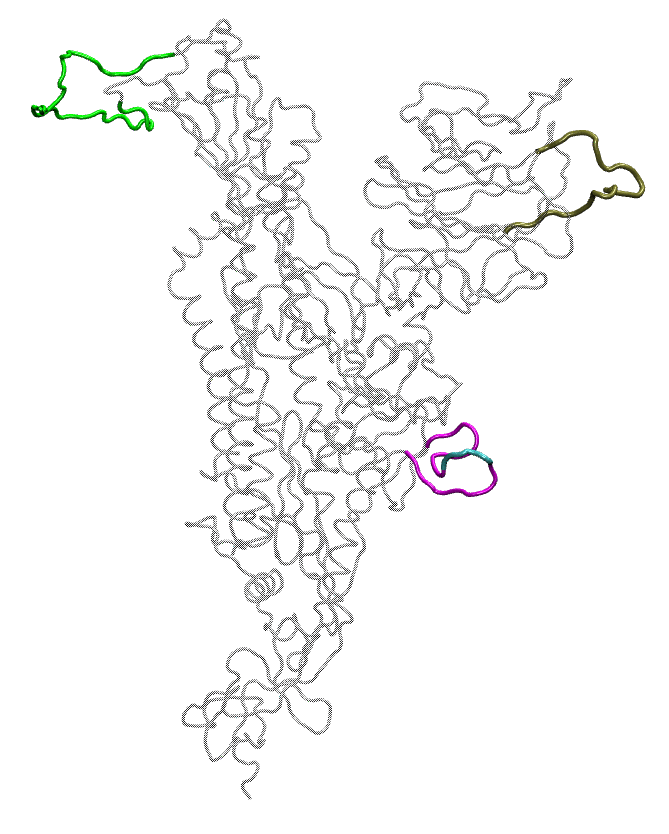
